# A non-invasive method to genotype cephalopod sex by quantitative PCR

**DOI:** 10.1101/2025.10.28.685099

**Authors:** Frederick A. Rubino, Gabrielle C. Coffing, Connor J. Gibbons, Scott T. Small, Thomas Desvignes, Jeffery Pessutti, Ann M. Petersen, Alexander Arkhipkin, Zhanna Shcherbich, John H. Postlethwait, Andrew D. Kern, Tessa G. Montague

## Abstract

Coleoid cephalopods (cuttlefish, octopus, and squid) are emerging model organisms in neuroscience, development, and evolutionary biology, and are of major economic importance in global fisheries. However, they are notoriously difficult and expensive to culture. The ability to determine sex early in development would enable more efficient and sustainable population management in both laboratory and wild settings. Here, we present a non-invasive method to genotype the sex of dwarf cuttlefish (*Ascarosepion bandense*) as young as three hours post-hatching using a skin swab and quantitative PCR assay, which detects a two-fold dosage difference between ZZ and Z0 sex chromosomes of males and females, respectively. Furthermore, we designed and validated primers for four additional cephalopod research species with assembled genomes (*Octopus bimaculoides, Sepia officinalis, Euprymna berryi*, *Doryteuthis pealeii*), and for a wild-caught species of economic value (*Illex illecebrosus*) using low-coverage whole genome sequencing data. This method enables accurate sex determination from hatchlings to adults across cephalopods, independent of genome quality or availability.

**Highlights:**

- The Z sex chromosome was identified in multiple cuttlefish, squid, and octopus species.
- A sensitive quantitative PCR assay can genotype ZZ/Z0 sex in each species.
- Low-coverage short-read sequencing data is sufficient to design effective primers.
- qPCR on non-invasive skin swabs enables genotyping of living animals.

## Introduction

The coleoid cephalopods – octopus, cuttlefish and squid – are a group of marine mollusks with a striking array of biological adaptations, including dynamic camouflage, regenerative limbs, the largest brain-to-body-ratio of the invertebrates, tactile chemosensation (“taste by touch”), extensive RNA editing, and skin-based social communication^1–6^. The last common ancestor of the cephalopods and vertebrates lived ∼600 million years ago^7^, prior to the emergence of centralized nervous systems. Since then, cephalopods have evolved the largest and most complex brains among invertebrates – capable of learning, memory, problem-solving and adaptive camouflage^8–13^. There is growing interest in the scientific study of cephalopod biology, which has been bolstered by the recent development of cephalopod tools, including genome and transcriptome sequences^14–18^, brain atlases^19–23^, cell atlases^24–27^, transient and multigenerational mutant animals^28, 29^, developmental staging series^30–32^, computational behavioral tools^33–36^, neural imaging using calcium dyes^37^, and electrophysiological recordings^38–40^.

Cephalopod research has traditionally been conducted in marine stations, relying on seasonal access to wild-caught cephalopods^41^. Over the past decades, however, multiple species have been successfully cultured through multiple generations in closed aquatic systems. Prominent species include the dwarf cuttlefish (*Ascarosepion bandense,* formerly *Sepia bandensis*)^30^, California two-spot octopus (*Octopus bimaculoides*)^42^, common cuttlefish (*Sepia officinalis*)^43^, Hawaiian bobtail squid (*Euprymna scolopes*)^44^, hummingbird bobtail squid (*Euprymna berryi*)^45^, and flamboyant cuttlefish (*Ascarosepion pfefferi*, formerly *Metasepia pfefferi*)^46^. These multi-generational cultures can provide year-round access to freshly fertilized eggs for microinjection, embryos of any stage for developmental studies, individuals at defined life stages for behavioral assays, and the opportunity to establish mutant or transgenic lines. Despite these advantages, stable and cost-effective cephalopod culture remains one of the greatest barriers to scientific progress: Cephalopods have high metabolic demands, are extremely sensitive to water quality, and have complex behaviors^47^. Thus, running a cephalopod facility requires specialist cephalopod expertise and considerable research expenses in equipment, personnel, and food costs.

Sex ratios in captive cephalopod populations can have a major impact on survival, behavior, and egg yield. For some cuttlefish species, mixed-sex groups can be maintained safely, but an excess of males can reduce egg production and increase aggression^48^. In many other cuttlefish species, mixed sex groups can become highly aggressive at sexual maturity, with males sometimes cannibalizing conspecifics unless housed in large holdings or kept in isolation^49^. By contrast, groups composed entirely of males can exhibit markedly lower aggression (unpublished observation). Sex-based management strategies can therefore be successful, but sex must be determined prior to sexual maturity. Anatomical determination of sex is theoretically possible: most mature male coleoid cephalopods possess a hectocotylus – a modified arm with fewer suckers, which is used to transfer spermatophores^13^ – but this modification can be subtle and difficult to detect. Some species exhibit sex-specific body sizes or social skin patterns, but such differences typically appear only after sexual maturity^13^, limiting their utility for early-stage sex identification. Beyond husbandry, accurate sex identification is essential for behavioral research. During social interactions, cuttlefish use a repertoire of innate skin patterns to communicate^6^, some of which are sex-specific. However, some species have been observed mimicking the opposite sex through skin patterning^50, 51^, making behavioral cues insufficient for reliable identification. A non-invasive method to determine sex in very young cephalopods could enable researchers to establish optimal sex ratios, which could in turn reduce aggression, increase reproductive output, and improve animal welfare. Additionally, an optimized sex ratio could reduce the number of animals required in a facility, potentially saving many thousands of dollars in yearly research costs and facilitating more rigorous experimental design.

The recent chromosome-level assembly of the *Octopus bimaculoides* genome revealed the presence of a genetic sex determination system in cephalopods, consisting of a Z0 sex karyotype in females and ZZ sex karyotype in males^52^. With this system, sex could be determined by measuring the relative abundance of Z-linked sequences, which are present at twice the level in males compared to females. This principle was recently used to determine sex in bigfin reef squid (*Sepioteuthis lessoniana*)^53^. Here, we identified the orthologous Z chromosome in *A. bandense*, *S. officinalis*, *E. berryi*, and *D. pealeii*, optimized a method to non-invasively extract genomic DNA from live cephalopods as young as three hours post-hatching (8 mm body length; 5.3 mm mantle length *A. bandense*), and established a sensitive quantitative PCR (qPCR) protocol that can detect a two-fold difference in the number of Z sex chromosomes present in a DNA sample relative to autosomes, thereby differentiating males from females (Figure 1). Our method was 100% accurate, based on the confirmed post-mortem analysis of 81 *A. bandense*. Furthermore, we designed and validated qPCR primers for *O. bimaculoides, S. officinalis*, *E. berryi*, and *D. pealeii,* and identified Z-linked genes and primers in *Illex illecebrosus* using only low-coverage short-read data. The *I. illecebrosus* primers also accurately distinguished sex in the related *Illex argentinus*, demonstrating that this approach can be extended to species without a complete genome assembly. This method promises to facilitate the use of cephalopods in laboratory culture and research.

**Figure 1.**
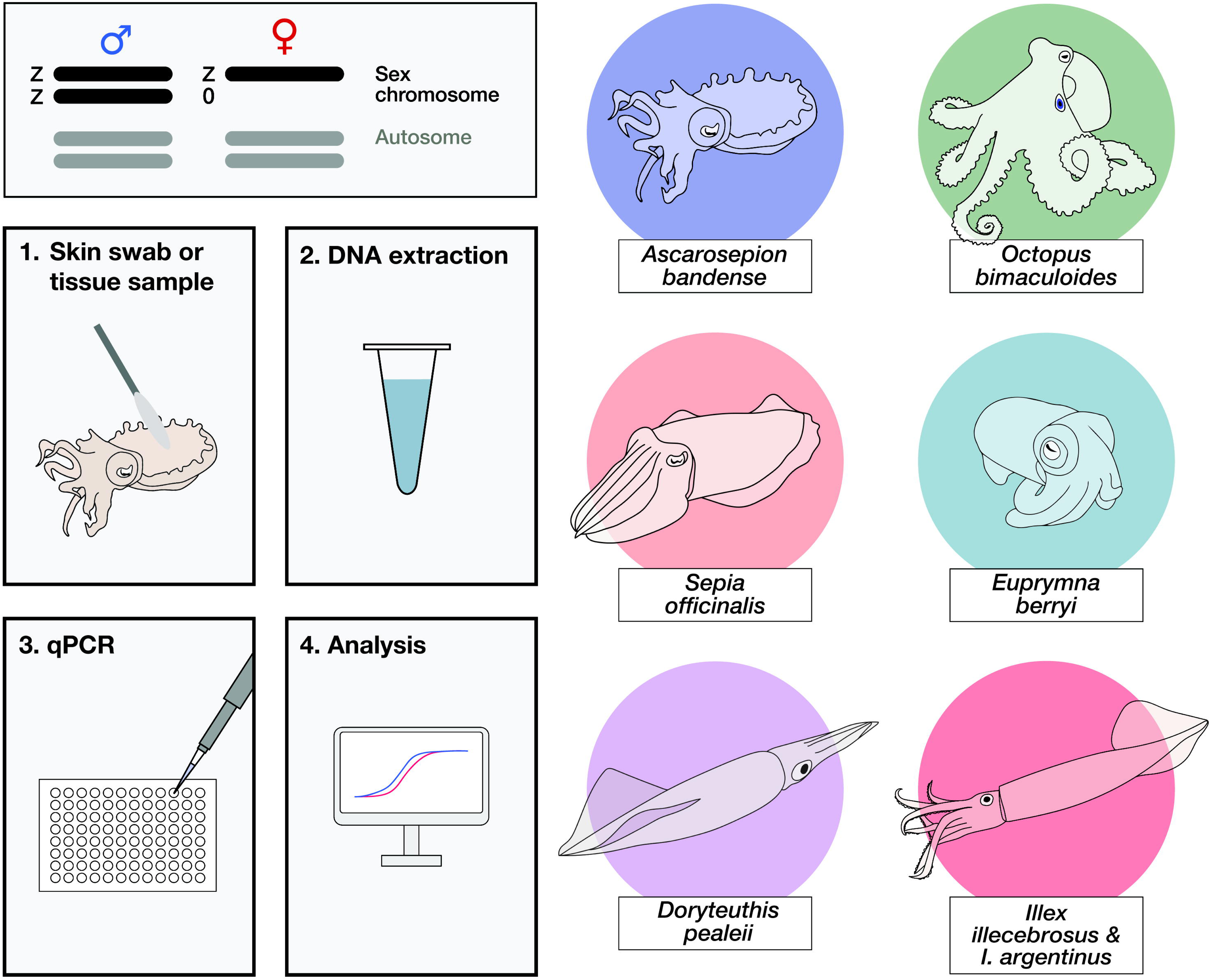
Overview of the sex genotyping protocol. Coleoid cephalopods, including dwarf cuttlefish (*Ascarosepion bandense*), California two-spot octopus (*Octopus bimaculoides*), common cuttlefish (*Sepia officinalis*), hummingbird bobtail squid (*Euprymna berryi*), longfin inshore squid (*Doryteuthis pealeii*), northern shortfin squid (*Illex illecebrosus*), and Argentine shortfin squid (*Illex argentinus*), appear to have a ZZ/Z0 sex determination system, in which males carry two copies of the Z chromosome and females carry one. Sex is determined by measuring the abundance of a Z-linked sequence relative to an autosomal sequence using qPCR after genomic DNA extraction from a skin swab or tissue sample.

## Results

### Conserved synteny and homology reveal orthologous sex chromosomes in multiple coleoid species

A cephalopod sex chromosome (Z) was previously identified within the California two-spot octopus (*Octopus bimaculoides*) genome (chromosome 17)^52^. To identify the putative Z chromosomes in other species, we examined conserved synteny across chromosome-scale genome assemblies of six coleoid cephalopods: *Sepia esculenta*^17^, *S. officinalis*^17^, *E. berryi*^24^, *E. scolopes*^15^, *D. pealeii*^14^, and *O. bimaculoides*^52^. In *S. esculenta* and *E. scolopes*, the Z chromosomes were identified previously^52^. In three additional species, we identified the putative Z chromosomes: chromosome 43 (*D. pealeii*), chromosome 44 (*E. berryi*), and chromosome 49 (*S. officinalis*, consistent with other findings^54^) (Figure 2, Figure S1).

**Figure 2.**
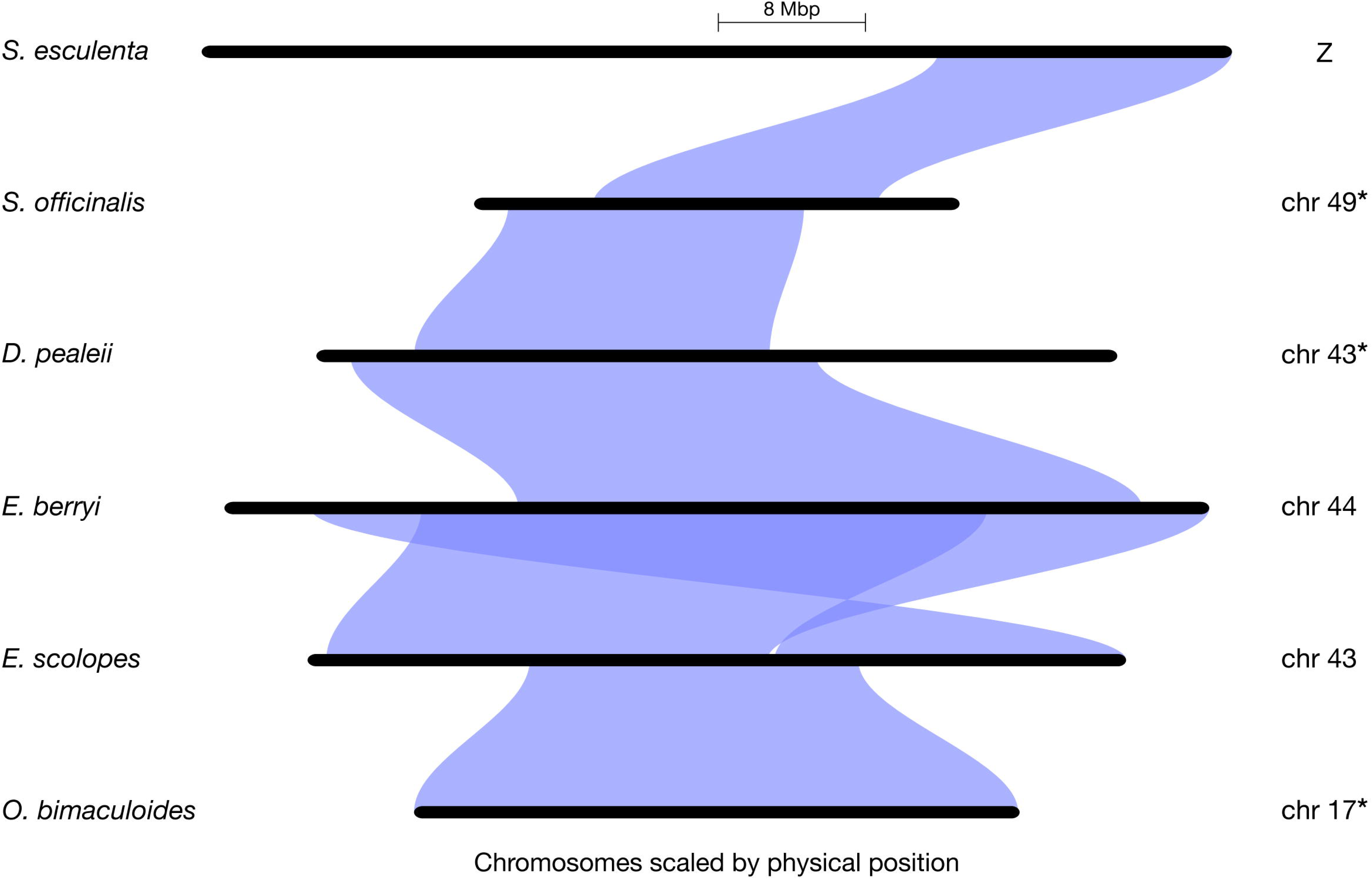
Conserved synteny and homology reveal orthologous sex chromosomes in multiple coleoid species. Riparian synteny plot of the Z chromosomes of *O. bimaculoides, E. scolopes*, *E. berryi*, *D. pealeii*, *S. officinalis*, and *S. esculenta,* with *E. scolopes* set as the reference species. Z chromosomes from *S. officinalis*, *D. pealeii*, and *O. bimaculoides* (marked with asterisks) were inverted to improve visualization. See also Figure S1.

The current *A. bandense* genome assembly^55^ is not at chromosome level, so we could not include it in our conserved synteny analyses. Therefore, to identify the putative Z contigs in this species, we aligned the *A. bandense* genome scaffolds to the *E. scolopes* genome and extracted contigs that aligned to the *E. scolopes* Z chromosome (chromosome 43, see Methods).

### Quantitative PCR shows sex-specific dosage differences in octopus, cuttlefish, and squid species

To test whether qPCR could be used to determine the sex of *A. bandense, O. bimaculoides, S. officinalis*, *E. berryi,* and *D. pealeii*, we designed primer sets targeting the Z chromosome (“sex” primers) and an autosome (“autosome” primers) using the Z chromosome sequences we identified (Figure 2). The goal was to measure whether the sex amplicon amplifies at equal the dosage (male) or half the dosage (female) of the autosome control. A two-fold difference in DNA template dosage amounts to a one-cycle difference in amplification; thus, we required primers with high specificity that generate a single melt curve representing a single product. Using stringent parameters (see Methods), we designed 24 primer sets for each species and tested them on DNA extracted from arm tissue from a confirmed phenotypic male and a confirmed phenotypic female cuttlefish. At least one primer set for each species yielded a ΔΔC_q_ difference of 1 between the male and female DNA samples (see Methods), demonstrating sufficient sensitivity to detect a two-fold difference in sex chromosome dosage and generating a single amplicon (Table 1, Figure S2, Table S1). Using blinded, post-mortem tissue from eight individuals of known phenotypic sex per species, we performed qPCR and examined the resulting ΔΔC_q_ values. In every case, the values segregated into two clusters corresponding to males (ΔΔC_q_ = 0) and females (ΔΔC_q_ = −1), yielding 100% correct assignments in each species (Figure 3A-E, Figure S3, 8/8, *p* = 0.0039, one-sided exact binomial test). Mean values of the male and female groups for each species were compared using Student’s t-tests (unpaired two-sample for all species except *O. bimaculoides*, for which a one-sample t-test was used) and were significantly different in all cases (*p* < 10^−4^; see Methods).

**Figure 3.**
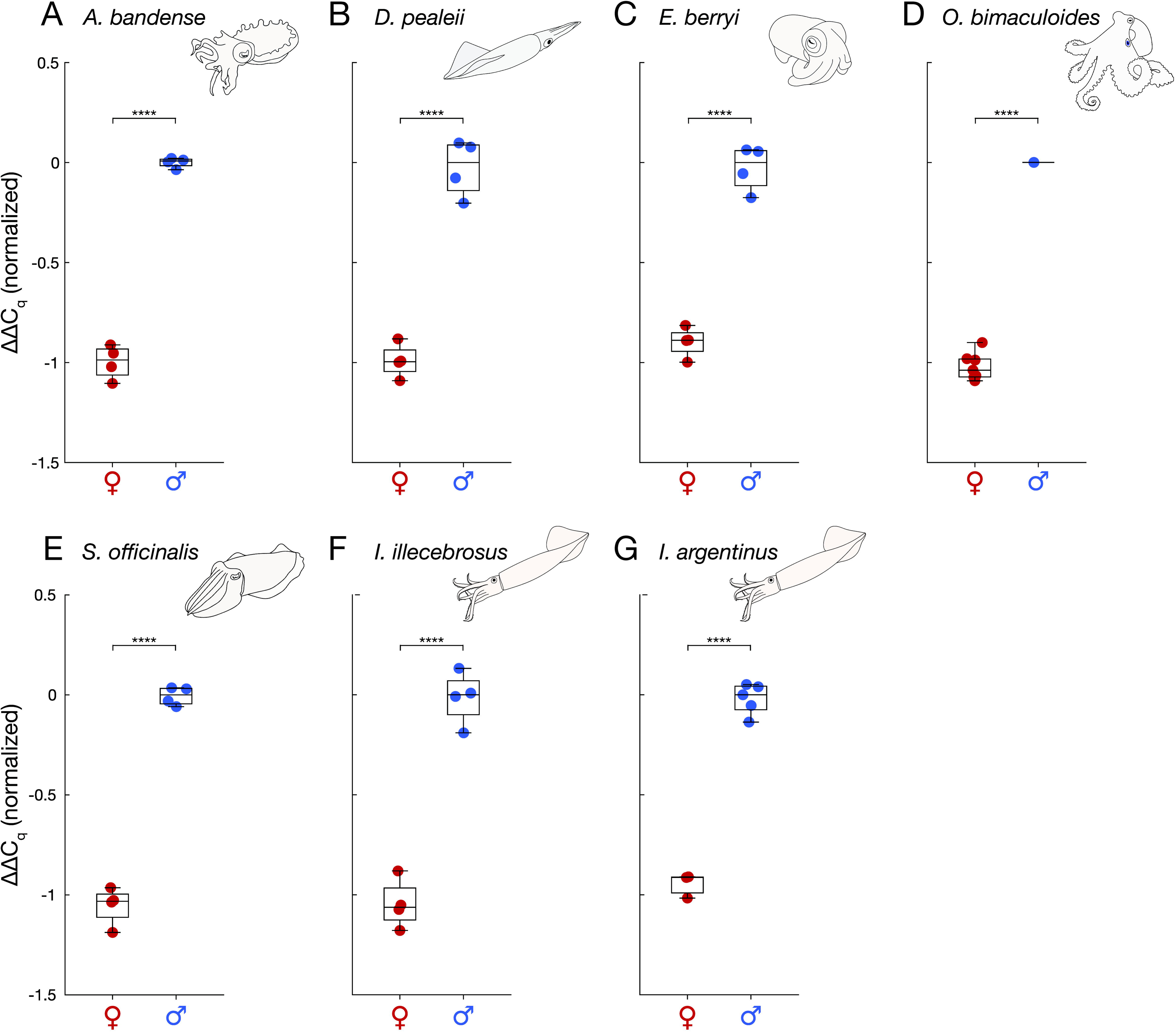
Quantitative PCR shows sex-specific dosage differences in seven octopus, cuttlefish, and squid species. (A) *A. bandense.* (B) *D. pealeii.* (C) *E. berryi.* (D) *O. bimaculoides.* (E) *S. officinalis.* (F) *I. illecebrosus.* (G) *I. argentinus.* For each species, ΔΔC_q_ values were calculated between autosomal and sex-linked loci across eight blinded, randomized tissue samples. Values were normalized such that the median of the higher-ΔΔC_q_ cluster (male) was set to zero. Each point represents the mean value of four technical replicates from a single animal (N = 8 animals for each species). Datapoints are colored by post-mortem-confirmed sex (females, red; males, blue). **** *p* < 1 × 10⁻⁴. See also Figures S2-S4.

**Table 1.**
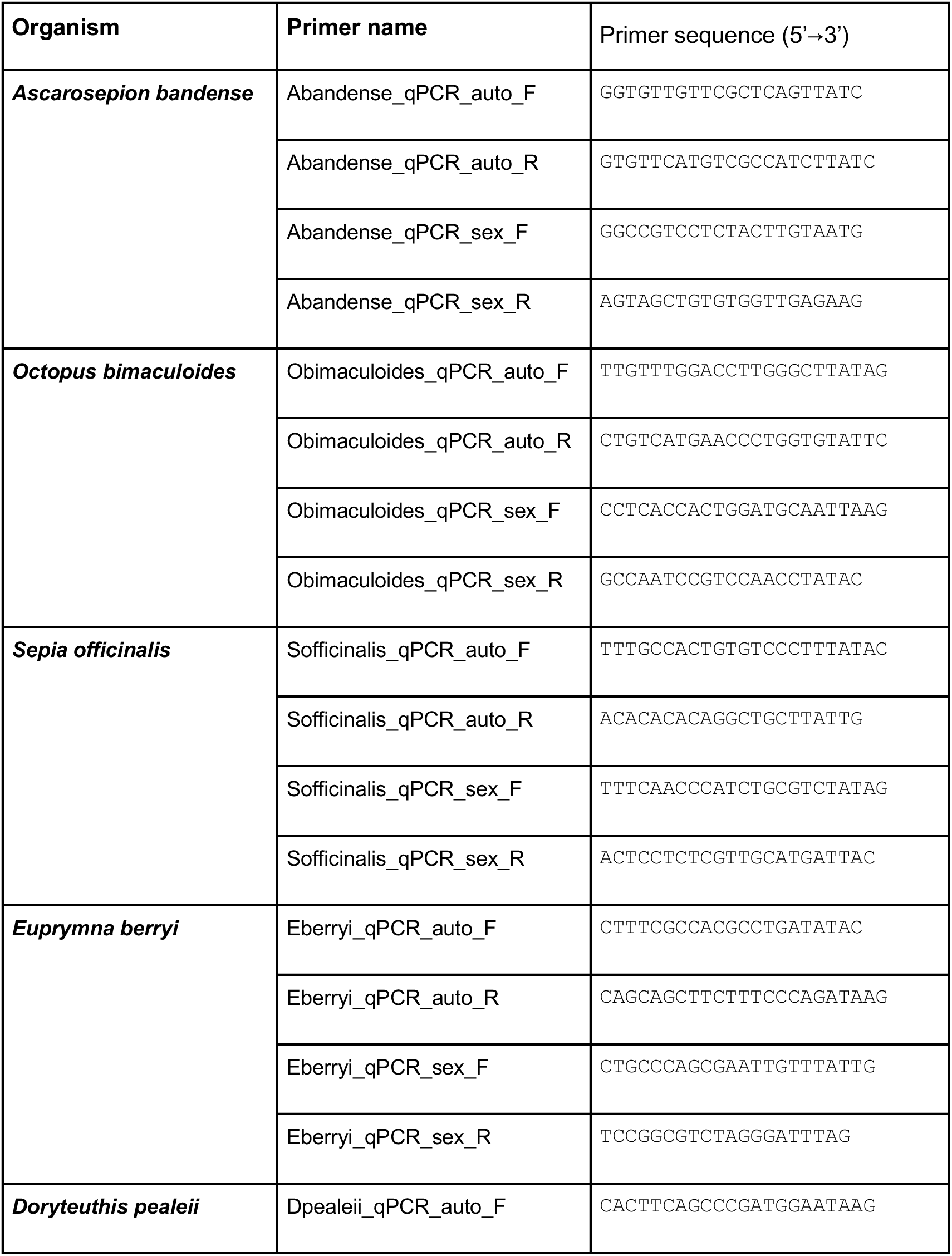
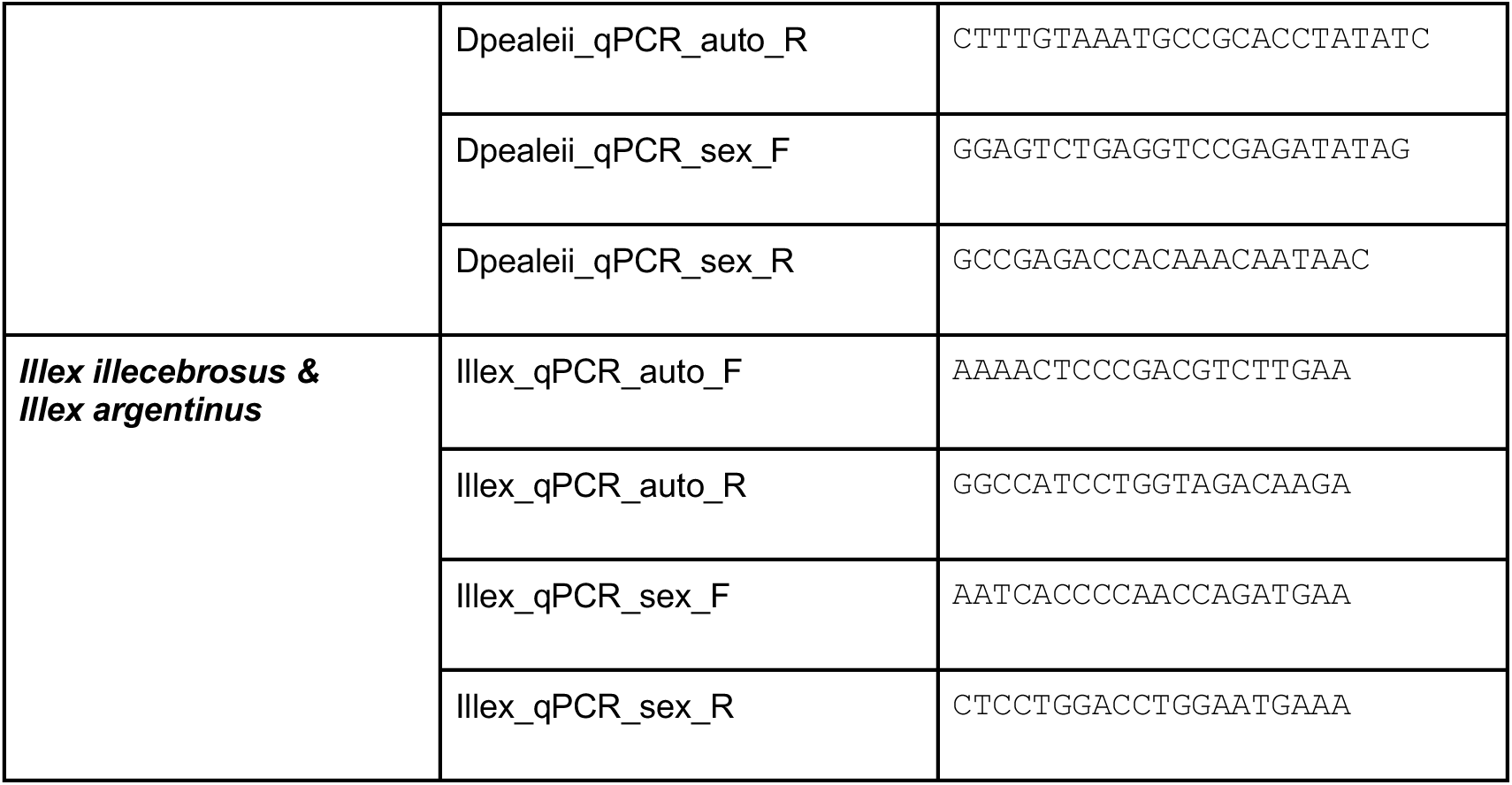
qPCR primer sequences for cuttlefish, octopus and squid species. For additional primer details, see Table S1.

Many cephalopod species of biological and economic interest, such as the squid *lllex illecebrosus,* lack chromosome-level genome assemblies. To test whether short-read sequencing data are sufficient for primer design, we generated low-coverage whole genome sequencing data from *I. illecebrosus* and aligned them to the genome of the most closely related species with a reference genome, *D. pealeii*. Based on *D. pealeii* sequences, we designed a panel of 24 primer sets (12 sex, 12 autosome) and then refined the primer sequences to match *I. illecebrosus*-specific variants (Figure S4). From this panel, one primer set displayed the required performance (Table 1, Figure S2). We validated the primers on eight blinded *I. illecebrosus* individuals of known phenotypic sex, and achieved 100% sex attribution accuracy (Figure 3F, 8/8, *p* = 0.0039, one-sided exact binomial test). Notably, the same primers also correctly identified sex in eight blinded samples of the related Argentine shortfin squid *Illex argentinus* (Figure 3G, 100%, 8/8, *p* = 0.0039). Thus, qPCR targets can be designed from low-coverage short-read sequencing data, and at least in some cases, can be functional for related species that lack extensive genome information.

### Quantitative PCR from non-invasive skin swabs successfully identifies sex

A qPCR assay to determine animal sex would be most useful if it could be applied to live animals. Cutaneous swabs have been successfully coupled with qPCR for clinical diagnosis^56^ and animal pathogen detection^57^. We therefore adapted a previously described squid genotyping protocol^29^ to swab the mantle skin of live, unanesthetized *A. bandense* and extracted DNA from the swabs (100% survival, n = 81, Figure 4A,B). After running a qPCR with the validated *A. bandense* autosome and sex primers (Figure 3A), results yielded ΔΔC_q_ values that clustered into two groups (median value distance of 0.97 and separation of 0.44 between the nearest data points from opposite groups) corresponding to putative males and females (Figure 4C, Figure S5A). To validate these assignments, we performed post-mortem dissections 3 months later after the animals reached sexual maturity, which revealed that the qPCR results were 100% accurate (81/81 correct, *p* = 4.14×10^−25^, one-sided exact binomial test; Figure 4C, Figure S5A). To find out how early in development animals could be swabbed and successfully sex genotyped, we swabbed 20 hatchlings 3 hours post-hatching (5.3 mm mantle length; Figure 4D) and raised them for 2 weeks to ensure their survival. All swabs yielded sufficient DNA for amplification by qPCR, all animals survived (20/20), and the genotyping results matched those obtained from post-mortem tissue extractions (Figure S5B-D). Together, these results establish that a non-invasive skin swab can provide sufficient DNA for qPCR-based sex genotyping in *A. bandense*, even from newly hatched animals as small as ∼5 mm mantle length.

**Figure 4.**
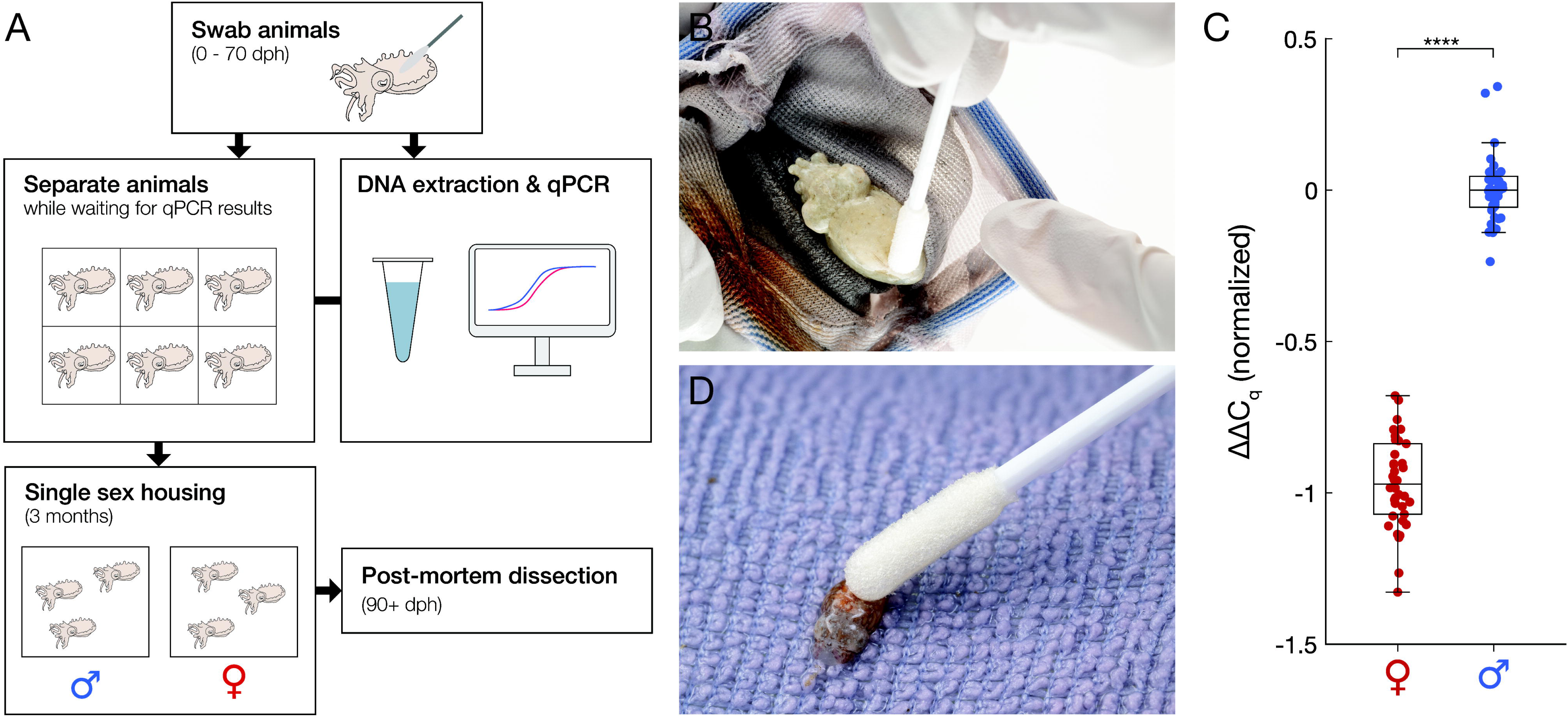
Quantitative PCR from non-invasive skin swabs successfully identifies sex. (A) Live animal sex genotyping workflow. Juvenile animals were swabbed and individually housed during the DNA extraction and qPCR. After obtaining qPCR results, animals were group housed by predicted genetic sex until confirmation of phenotypic sex by dissection. dph, days post-hatching. (B) A 10-week-old juvenile being swabbed for DNA extraction. (C) ΔΔC_q_ values between autosomal and sex loci for 81 live juvenile *A. bandense* individuals, normalized such that the median value of the male cluster is set to ΔΔC_q_ = 0. Each data point represents the mean value of four technical replicates from a single animal (N = 81 animals). Predicted sex is indicated as red (female) or blue (male). All 81 predictions were confirmed by post-mortem dissection. **** *p* < 1 × 10⁻⁴. (D) A 3-hour-old hatchling being swabbed for DNA extraction. See also Figure S5.

## Discussion

This study demonstrates that a non-invasive skin swab combined with a qPCR assay can reliably determine the sex of coleoid cephalopods. The workflow is efficient: in our hands, swabbing 48 animals, extracting DNA, setting up and running a qPCR, and analyzing the results can all be completed within a single day (Figure 1). Importantly, this approach enables accurate sex determination across life stages – from hatchlings to adults – and across a range of cephalopod species, independent of genome quality or availability. Sex genotyping by qPCR can also be applied to embryos, and to post-mortem tissue, as has been demonstrated in the bigfin reef squid^53^ and here (Figure 3).

Recent genomic analyses of Nautilus, a highly divergent cephalopod clade, suggest the presence of an XX/XY sex determination system^58^. This raises the possibility that additional cephalopod lineages may have independently evolved alternative mechanisms of sex determination. Although the methodological approach presented here could be applied to Nautilus to detect a two-fold X-chromosome dosage difference between males and females, a simpler approach would be to use standard PCR with primers targeting sequences on the Y chromosome.

This method has the potential to have a major impact on captive cephalopod population management. Populating breeding tanks with optimal sex ratios can maximize egg production while minimizing aggression. In turn, this can reduce the number of animals required in a culturing facility, reducing operational costs. The nature of this procedure allows for quick and efficient DNA collection, enabling accurate sex identification without the need for anesthesia or any invasive methods. These goals are in line with the principles of the 3Rs (Replacement, Reduction, and Refinement)^59^ and can help guide future welfare considerations for cephalopods.

Creating breeding groups for genetic diversity or genetically-modified lines can be both inefficient and high risk when using traditional sex identification, such as behavioral observations or hectocotylus identification. Our unpublished observations have shown that female *A. bandense* that are paired with males after they reach sexual maturity lay fewer eggs and are more susceptible to aggression or even cannibalism by males than those paired before sexual maturity. Furthermore, female cephalopods can carry sperm from different mates^60^ for days to months^61^, making any females that have previously lived with sexually mature males poor candidates for selective breeding. The ability to determine sex prior to sexual maturity permits longer cohabitation, which can reduce aggression and eliminate the risk of undesired sperm storage in females – a crucial resource in the pursuit of generating stable transgenic or mutant lineages.

Beyond husbandry applications, non-invasive sex genotyping could facilitate sex identification in juvenile cephalopods used in growth or physiological studies, identify environmentally mediated sex reversals^62^, and support fisheries management. For example, the northern and Argentine shortfin squids, *I. illecebrosus* and *I. argentinus,* are economically important species that are harvested in large volumes for seafood markets. The ability to non-destructively determine the sex of live animals, especially paralarvae or juveniles, could enable the assessment of sex ratios in wild populations during monitoring surveys, and assist fisheries managers and law enforcement in promoting sustainable harvesting through sex-specific catch limits. For such applications, portable rapid field assays would be more practical than laboratory-based qPCR, highlighting an important area for future development.

## Limitations of the study

There are two main limitations that impact the practicality of this method. First, qPCR instruments are expensive, and not all research laboratories or fisheries research vessels will have routine access to one. One solution could be to use commercial providers that offer fee-for-service qPCR, although costs and turnaround times may limit feasibility, especially given the need to house animals individually after swabbing. Given that standard qPCR is not inherently compatible with field deployment, the assay could in principle be adapted to isothermal amplification approaches that operate at a single temperature and permit simple visual or lateral-flow readouts, such as loop-mediated isothermal amplification (LAMP) assays^63^, which can be quantitative^64, 65^ or recombinase polymerase amplification (RPA) assays^66^. Such adaptations would substantially broaden usability outside the laboratory.

Second, primer design requires some genetic sequence information for the target species. However, as our results show, low-coverage whole genome sequencing reads can be aligned to a closely related species’ genome assembly to identify conserved loci in autosomes and sex chromosomes. As additional cephalopod genomes become available, it may become possible to design universal primers that function across multiple species, expanding accessibility, although deeper sequence divergence is likely to limit truly universal primer design.

In summary, non-invasive sex genotyping by qPCR provides a versatile and accurate tool for research, enabling improved breeding, animal welfare, and resource management across cephalopods.

## Resource availability

### Lead contact

Requests for further information, resources, and reagents should be directed to and will be fulfilled by the lead contact, Tessa Montague.

### Materials availability

All items described here are commercially available.

## Acknowledgements

We would like to thank Richard Axel for valuable discussions, funding, and mentorship; Ruth Lehmann for funding and mentorship; Gjendine Voss (Rosenthal lab), Kendra Buresch (Hanlon lab), Lisa Abbo, Bret Grasse, Saurabh Attarde (Holford lab), and Catherine Wilson for tissue samples; Gabrielle Winters Bostwick and Sarah Detmering (Crook lab) for testing the protocol and providing feedback; Silas Tittes for valuable discussions; Dan Rokhsar for sharing the unpublished *E. berryi* genome; Josh Rosenthal for sharing additional *E. berryi* sequencing data; Thomas Barlow for photographing the swabbing process; and Thomas Mungioli for designing and constructing the cuttlefish holding system that allowed animals to be separated for large-scale genotyping trials. We also thank Jessica Izguerra (former NOAA intern) and Sarah Salois (former NOAA Fisheries biologist) for training on *Illex* sex ID and measurements, and Captain Stefan Axelsson and Chief Engineer David Axelsson of the FV Dyrsten for their assistance in sample collection, willingness to provide their catch, and for allowing J.P. to sail on their vessel. This project reflects NOAA’s Cooperative Research model of scientists working collaboratively with industry and would not have been possible without the relationships formed through this approach. F.A.R. was supported by the Whitehead Institute (through Dr. Ruth Lehmann); C.J.G., and T.G.M. were supported by the Howard Hughes Medical Institute (HHMI, through Dr. Richard Axel); G.C.C, S.T.S, S.T., and A.D.K. were funded in part through NIH Awards R01HG010774 and R35GM148253. NIH award R35GM139635 and NSF grant 2232891 to J.H.P and T.D. contributed to this work. F.A.R. is supported by the Helen Hay Whitney Foundation, and T.G.M. is supported by the HHMI Hanna H. Gray Fellowship.

## Author contributions

Conceptualization: C.J.G., G.C.C., A.D.K., T.G.M.; Methodology: F.A.R., G.C.C., C.J.G., S.T.S., T.G.M.; Investigation: F.A.R., C.J.G., T.D., T.G.M.; Formal analysis: F.A.R., G.C.C., S.T.S.; Resources: C.J.G., J.P., A.M.P., A.A., Z.S.; Writing – original draft: T.G.M.; Writing – review & editing: F.A.R., G.C.C., C.J.G., S.T.S., T.D., J.H.P., A.D.K., T.G.M.; Visualization: F.A.R., G.C.C., T.G.M.; Project administration: A.D.K., T.G.M.; Supervision: J.H.P., A.D.K., T.G.M.

## Declaration of generative AI and AI-assisted technologies in the writing process

During the preparation of this work the authors used ChatGPT for light text editing. After using this tool, the authors reviewed and edited the content as needed and take full responsibility for the content of the published article.

## Materials and methods

### Key Resources Table

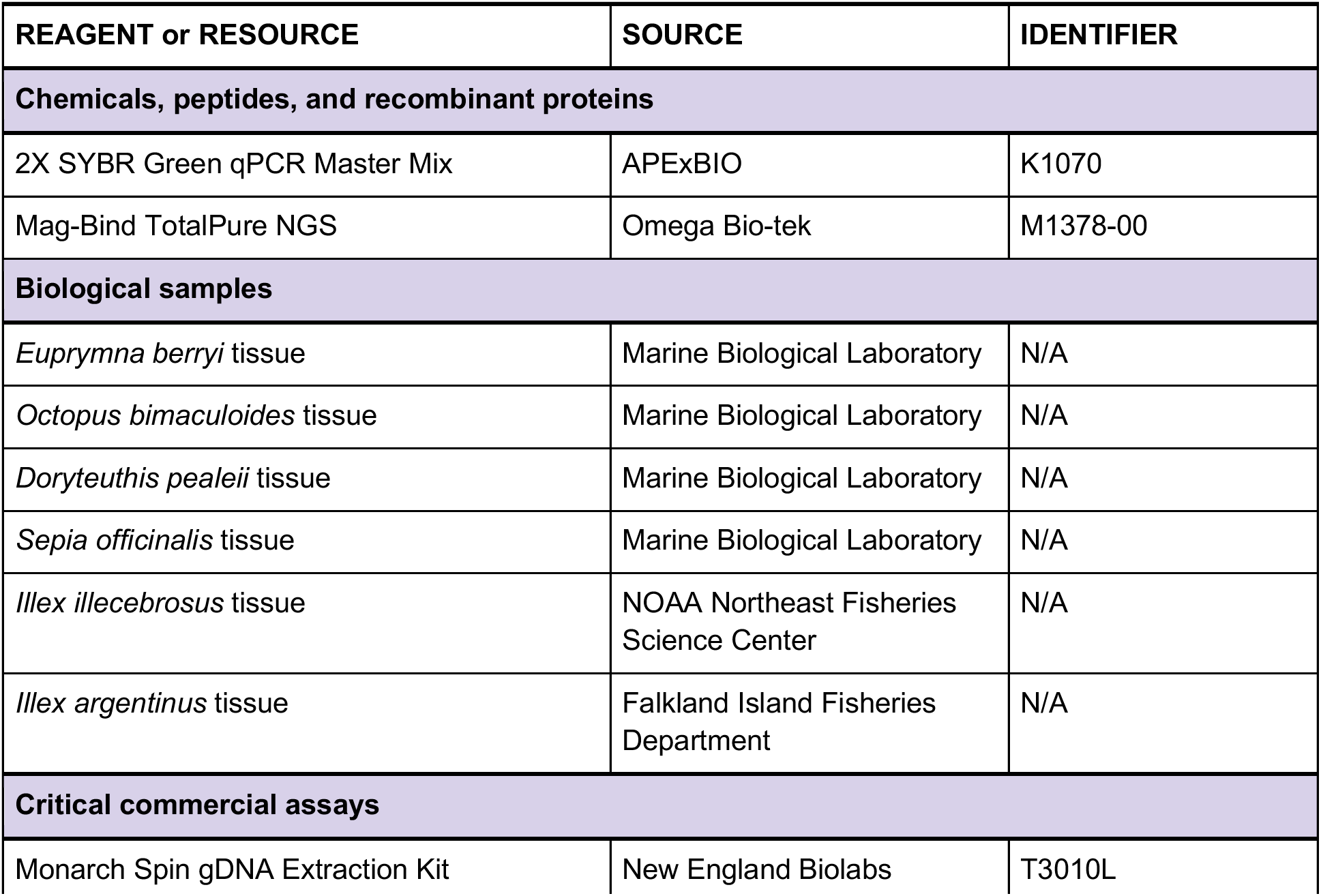
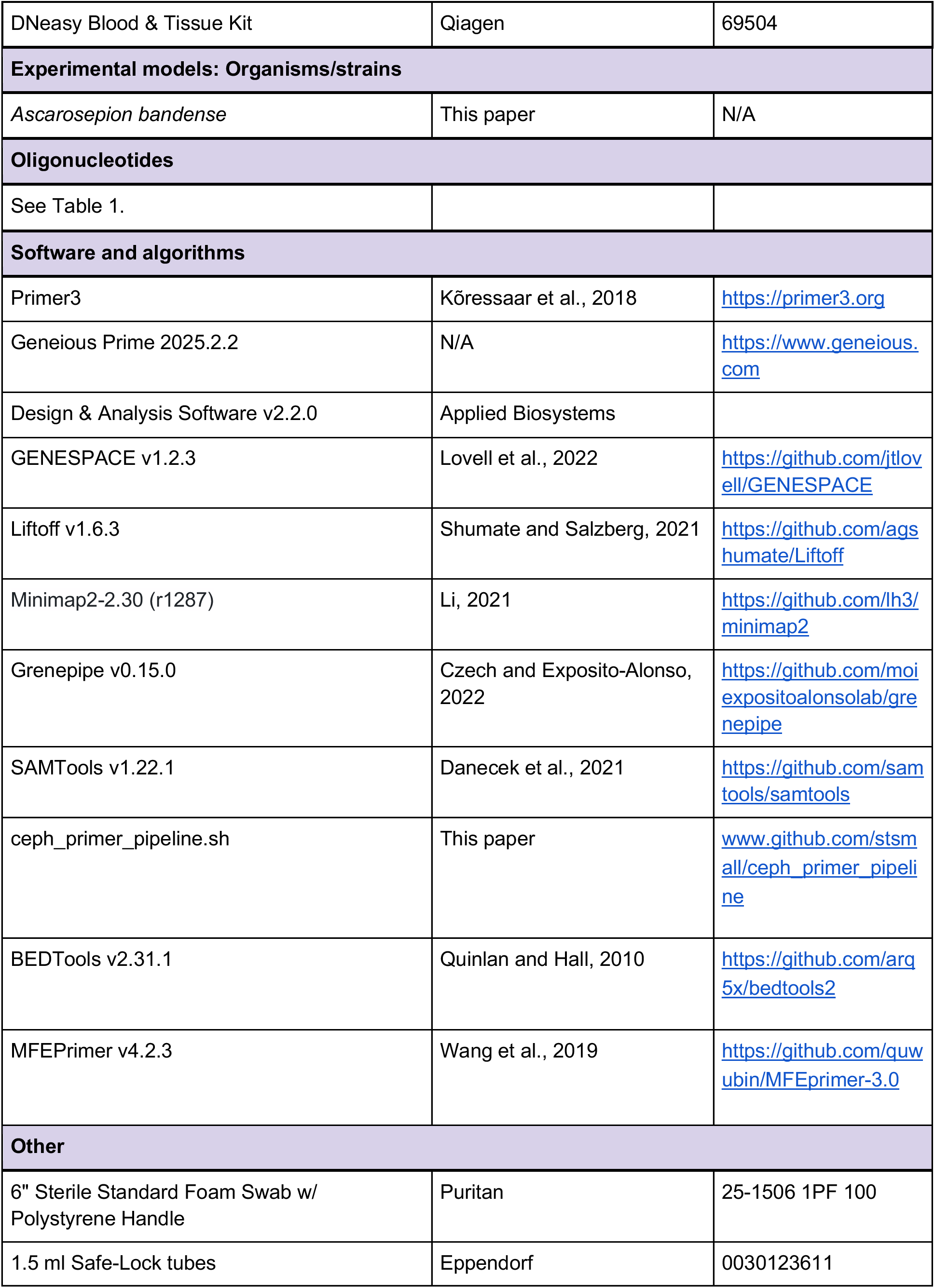
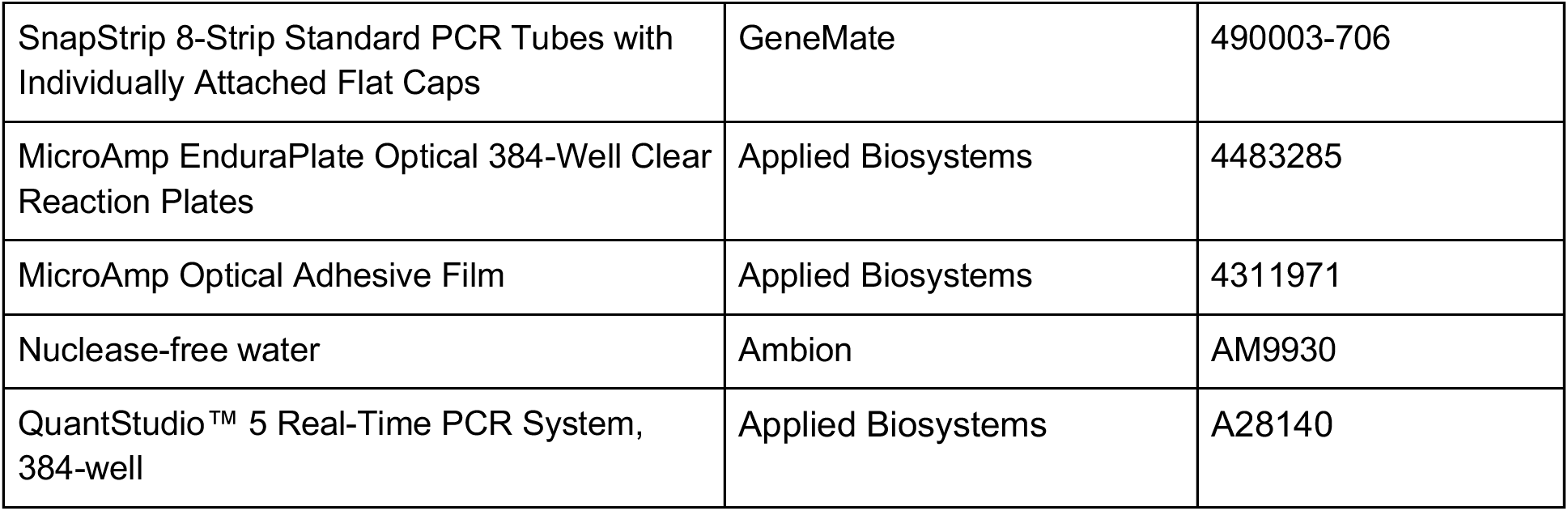

### Ethics statement

The use of cephalopods in laboratory research is currently not regulated in the USA. However, Columbia University has established strict policies for the ethical use of cephalopods, including operational oversight from the Institutional Animal Care and Use Committee (IACUC). All of the cuttlefish used in this study were handled according to an approved IACUC protocol (AC-AABT8673), including the use of deep anesthesia and the minimization and prevention of suffering.

### Z chromosome identification in *D. pealeii*, *E. berryi*, *S. officinalis,* and *A. bandense*

To conduct synteny analyses, peptide and gtf files for *D. pealeii* and *E. scolopes* were downloaded from their associated publications^14, 15^ while an unpublished assembly associated with the *E. berryi* cell atlas^24^ was used for *E. berryi* (see Table S1 for assembly information). An annotation was previously generated for *S. esculenta* by a gene annotation liftover from *S. pharaonis*^52^. At the time of our analyses, an annotation for the genome assembly of *S. officinalis* was not available. Therefore, we generated an annotation for *S. officinalis* by lifting over the genome annotation from *S. pharaonis* using Liftoff^67^ with default settings. Synteny analyses between the chromosomes of *S. esculenta*, *S. officinalis*, *E. berryi*, *E. scolopes*, *D. pealeii,* and *O. bimaculoides* were performed using the R package GENESPACE v.1.2.3^68^. In all synteny maps, *E. scolopes* was used as the reference species. In riparian plots and dotplots, the chromosomes are scaled by physical position (useOrder = False parameter in GENESPACE). The *A. bandense* genome assembly^55^ and associated transcriptome was pulled from cuttlebase.org/downloads. Since the *A. bandense* genome was not a chromosome level assembly, we could not use gene synteny to identify the putative Z chromosome. Instead, we used minimap2^69^ to align *A. bandense* to *E. scolopes* (with parameters -cx asm20) to identify potential Z contigs.

### Generation of *I. illecebrosus* short-read sequences and alignment to *D. pealeii* genome

To identify autosomal and sex-linked primers in *I. illecebrosus*, a set of primers was first designed for *D. pealeii* using orthologous genes identified by GENESPACE targeting chromosome Dpe01 (“autosome” primer) and chromosome Dpe43 (“sex” primer). Next, short-read sequencing data was generated from 20 *I. illecebrosus* individuals (10 males and 10 females, as determined by gonad morphology). Specifically, squid DNA was extracted using a DNeasy Blood & Tissue Kit (Qiagen) following the manufacturer’s instructions, and DNA extracts were purified and cleaned from short DNA fragments using TotalPure NGS Mag-Bind Beads (Omega Bio-tek). Low-coverage Whole Genome Sequencing Illumina libraries were prepared using the miniaturized Nextera DNA Library Preparation Kit with an Echo liquid handler and sequenced by the Genomics and Cell Characterization Core Facility (GC3F) at the University of Oregon using an Illumina NovaSeq 6000. Raw sequencing data was processed using grenepipe^70^ by aligning to the *D. pealeii* reference genome (GCA_023376005.1). The resulting BAM files were used to make a consensus using the SAMtools consensus tool^71^.

### qPCR primer design

For all species, 12 primer pairs were designed against autosomal genomic DNA sequences and 12 primer pairs against known or presumed sex chromosomal genomic DNA sequences. The targeted chromosome sequences are listed in Table S1. Primer design was performed as described below, and the final primer sequences, binding sites, and amplicon sequences are indicated in Table S1.

*A. bandense*, *O. bimaculoides*, *S. officinalis*, *E. berryi*, *D. pealeii*: Primers were designed using Geneious Prime 2025.2.2 (https://www.geneious.com) with a modified Primer3 (2.3.7,^72^) incorporating the following parameters: optimal primer size of 20 bp, optimal Tm of 59°C, optimal GC content of 50%, product size between 100-200 bp, GC clamp of 1, max poly-X of 3, and max dimer Tm 30°C. Primers were further screened for off-targets by a BLAST search against the appropriate reference genomes, and for secondary structures and self-complimentarity using IDT OligoAnalyzer (https://www.idtdna.com/calc/analyzer).

*I. argentinus*, *I. illecebrosus*: Primer pairs were generated using the ceph_primer_pipeline.sh (available at www.github.com/stsmall/ceph_primer_pipeline). This command-line script uses Primer3 to design primers (optimal primer size of 20 bp, optimal Tm of 60°C, optimal GC content of 50%, product size between 150-300 bp, and max poly-X of 5) and MFEprimer^73^ to check primer specificity against a reference genome. Using a GFF file and reference genome in FASTA format, this script uses BEDTools^74^ to extract user selected attributes (e.g. exons) from a given chromosome and builds a FASTA file for input into Primer3. The Primer3 default config file was modified to return only a single set of primers per FASTA entry, and the final primer sets are exported as a CSV file.

Primers were first tested against genomic DNA extracted from confirmed male and female individuals (see Genomic DNA extraction and Quantitative PCR sections below), as well as no-template controls. Primer pairs which gave a single, template-dependent melt curve were analyzed by 2% agarose gel electrophoresis and sequenced to confirm the generation of a single PCR product, which was validated against the reference sequence. The sequencing of PCR products was performed by Plasmidsaurus using Oxford Nanopore Technology with custom analysis and annotation (Premium PCR Sequencing service). Primer pair efficiency was measured by dilution series and confirmed to be within the accepted 90-110% range, in the concentration ranges used in this assay. For each species, a single autosome-targeting primer pair and a single sex chromosome-targeting primer pair were chosen for further sex genotyping analysis.

### Genomic DNA extraction from cephalopod frozen tissue

Frozen arm tips from *O. bimaculoides, S. officinalis, E. berryi,* and *D. pealeii* were obtained as flash frozen samples, while *I. illecebrosus* and *I. argentinus* samples were obtained as mantle fragments or arm tips preserved in 95% ethanol. All samples were temporarily stored at −80°C until DNA extraction. To extract genomic DNA, ∼3 mm pieces of tissue were placed in 1.5 ml Safe-Lock tubes, and processed using the Monarch Spin gDNA Extraction Kit. Briefly, 200 µl of Tissue Lysis Buffer and 10 µl of 20 mg/ml Proteinase K were added to each tube, and the samples were incubated at 56°C in a ThermoMixer shaking at 1400 rpm for 1-2 hours until fully digested. In some cases, the lysate turned pink or purple, due to chromatophore pigments. Following digestion, 400 µl of Binding Buffer was added, samples were pulse vortexed for 5-10 seconds, and the lysates were transferred to gDNA Purification Columns. Columns were centrifuged at 1,000 x *g* for 3 minutes followed by >12,000 x *g* for 1 minute, washed twice with 500 µl of gDNA Wash Buffer, and eluted with 60 µl of pre-warmed (60°C) gDNA Elution Buffer after a 5 minute incubation at room temperature. Eluted DNA concentrations ranged from ∼50 to 350 ng/µl. All DNA was diluted 20-fold (no further normalization required) and transferred into 8-well strips for subsequent qPCR.

### *A. bandense* husbandry

Dwarf cuttlefish (*Ascarosepion bandense*) were housed in a recirculating system of 30-35 ppt artificial seawater (Rea Sea Coral Pro Salt) maintained at 24-25°C. Water quality was maintained through frequent water changes and mechanical, chemical, and biological filtration with the following parameters: ammonia 0 ppm, nitrite <0.1 ppm, nitrate <20 ppm, and pH 8.0-8.4. Cuttlefish younger than 40 days post-hatching were fed live mysid shrimp (*Americamysis bahia*), after which they were transitioned to live grass shrimp (*Palaemonetes spp.*). Animals were fed three times daily.

### Skin swabbing and genomic DNA extraction from *A. bandense* juveniles and hatchlings

DNA extractions were performed using a Monarch Spin gDNA Extraction Kit with minor modifications to the manufacturer’s protocol, described below. Juvenile *A. bandense* were carefully caught with a soft net and positioned dorsal side up without anesthesia. A sterile foam swab was pre-soaked in tank water and gently stroked across the dorsal mantle skin from anterior to posterior, avoiding the anterior tip of the mantle (the most delicate region of the skin). As a practical note, swabbing adult *A. bandense* requires substantially greater pressure, likely due to the presence of mucus on the skin. The animal was then gently rotated ventral side up, and the other side of the swab was used to swab the ventral surface with slightly greater pressure. Each animal was placed in an individual tank to keep track of its identity, and the swab was placed face down into a 1.5 ml Safe-Lock tube preloaded with 200 µl Tissue Lysis Buffer and 3 µl of 20 mg/ml Proteinase K. After all samples were collected, swab shafts were trimmed to allow tube closure, and samples were incubated at 56°C in a ThermoMixer shaking at 1400 rpm for 1 hour. The shortened shaft of each swab was then held tightly while the swab was pressed and dragged against the side of the tube to recover liquid from the foam tip. Next, 400 µl of Binding Buffer was added, the samples were pulse vortexed for 5-10 seconds, and the lysate was transferred to gDNA Purification Columns. Columns were centrifuged at 1,000 x g for 3 minutes followed by >12,000 x *g* for 1 minute, washed twice with 500 µl of gDNA Wash Buffer, and eluted after a 5 minute incubation with 200 µl of pre-warmed (60°C) gDNA Elution Buffer. The DNA yield from an animal of 10-20 mm mantle length was typically ∼250 ng, although yields varied widely across samples. DNA was transferred into 8-well strips for qPCR, with no further normalization of concentration required.

To swab 3-hour-old hatchlings, a 5 L jug was filled with artificial seawater and covered with a soft microfiber towel such that the center of the towel rested approximately 5 mm below the water surface. Hatchlings were gently caught using a turkey baster and placed on top of the towel. To expose the mantle for swabbing, the sides of the towel were gently pulled upward, raising the animal so that the mantle was above the water surface (Figure S5C). A sterile foam swab pre-soaked in tank water was used to swab the mantle with extremely low pressure. The hatchling was then carefully rotated to a ventral-up orientation, and the other side of the swab was used to repeat on the ventral side. Each hatchling was placed in an individual tank to maintain identity until the skin swab qPCR results were acquired. The reminder of the protocol was the same as above, except that the DNA was eluted in a volume of 50 µl.

DNA yield from hatchling swabs was typically ∼13 ng, although yields varied widely across samples. DNA was transferred into 8-well strips for qPCR, with no further normalization of concentration required. After obtained the qPCR results for the hatchling swabs, the animals were grouped in tanks by predicted sex for two weeks to monitor post-swabbing survival. After two weeks, the hatchlings were euthanized in 10% ethanol in artificial seawater, a small piece of tissue was taken from each animal, and the “Genomic DNA extraction from cephalopod frozen tissue” protocol was followed.

### Quantitative PCR

qPCR reactions were performed on a QuantStudio 5 Real-Time PCR System (Applied Biosystems) using 2X SYBR Green qPCR Master Mix (APExBIO) with ROX reference dye (APExBIO). Reactions were prepared as technical repeats in quadruplicate (7 µL total volume, 0.25 µM forward primer, 0.25 µM reverse primer, 2 µL template DNA). Thermocycling conditions were as follows: initial denaturation (95°C for 2 min), amplification (95°C for 15 s, 60°C for 45 s, for a total of 45 cycles), melt curve analysis (95°C for 15 s, 60°C for 1 min, followed by an increase to 95°C at a rate of 0.06°C/s).

### Quantitative PCR analysis

C_q_ values, mean values, and standard error of the mean were automatically determined using Design & Analysis Software v2.2.0 (Applied Biosystems). Product identity was validated by melt curve comparison with sequence-verified standards (see qPCR primer design above). Outliers within a replicate group (C_q_ values differing by > 1 cycle from the mean) were discarded prior to analysis. ΔC_q_ values were calculated as C_q_ (autosome) minus C_q_ (sex chromosome) and were confirmed to cluster into two groups. ΔΔC_q_ values were calculated by subtracting the median value of the male (i.e., the greater ΔC_q_) cluster, or the value of a male standard. The midpoint ΔΔC_q_ value (−0.5) was used as the threshold between male and female classification. If samples produced a mean value within 1 standard error of the mean (SEM) of this value, they were considered insufficiently precise due to technical error and repeated. Reported SEM values for each individual represent the sum of its SEM values from autosome and sex replicate groups. Data visualization and analysis were performed in Design & Analysis Software, MATLAB R2024b, and Microsoft Excel. For each species, a one-sided exact binomial test was used to determine if the observed rate of correct predictions performed better than chance (50%). Mean ΔΔC_q_ values for male and female groups were compared using Student’s t-tests (unpaired two-sample tests for all species except *O. bimaculoides*, for which one-sample t-test against zero was used due to the availability of only a single male sample). In all cases, differences were highly significant (*p* < 1 × 10⁻⁴): *A. bandense*, *p* = 4.76 × 10⁻⁷; *D. pealeii*, *p* = 2.38 × 10⁻⁵; *E. berryi*, *p* = 1.37 × 10⁻⁵; *I. argentinus*, *p* = 2.13 × 10⁻⁶; *I. illecebrosus*, *p* = 2.77 × 10⁻⁵; *O. bimaculoides*, *p* = 2.06 × 10⁻⁸; *S. officinalis*, *p* = 1.06 × 10⁻⁶; and *A. bandense* juvenile skin swabs, *p* = 4.77 × 10⁻⁴⁹. Box-and-whisker plots show the median, interquartile range (25th–75th percentiles), and whiskers extending to the most extreme non-outlier values (±1.5×IQR).

