## Supplementary material for "A non-invasive method to genotype cephalopod sex by quantitative PCR": Figure S

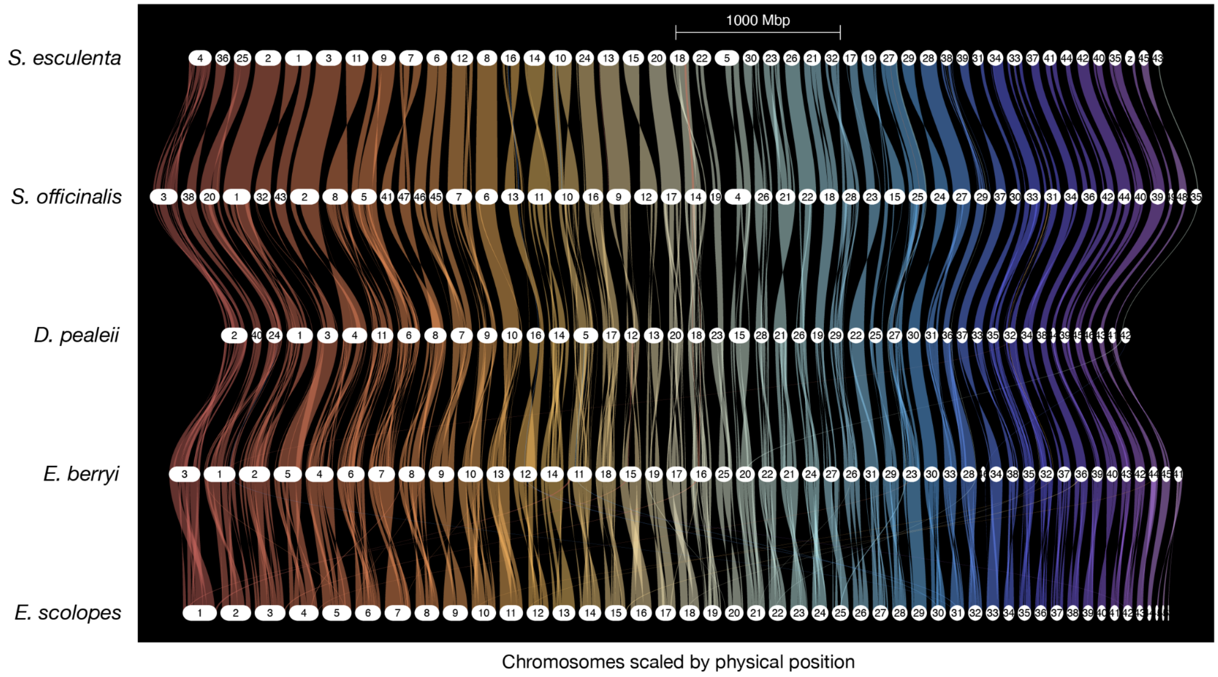


**Figure S1.** Conserved synteny relationships among chromosomes of *E. scolopes, E. berryi, D. pealeii, S. officinalis*, and *S. esculenta*. This riparian synteny plot was generated from orthogroups with *E. scolopes* set as the reference species.

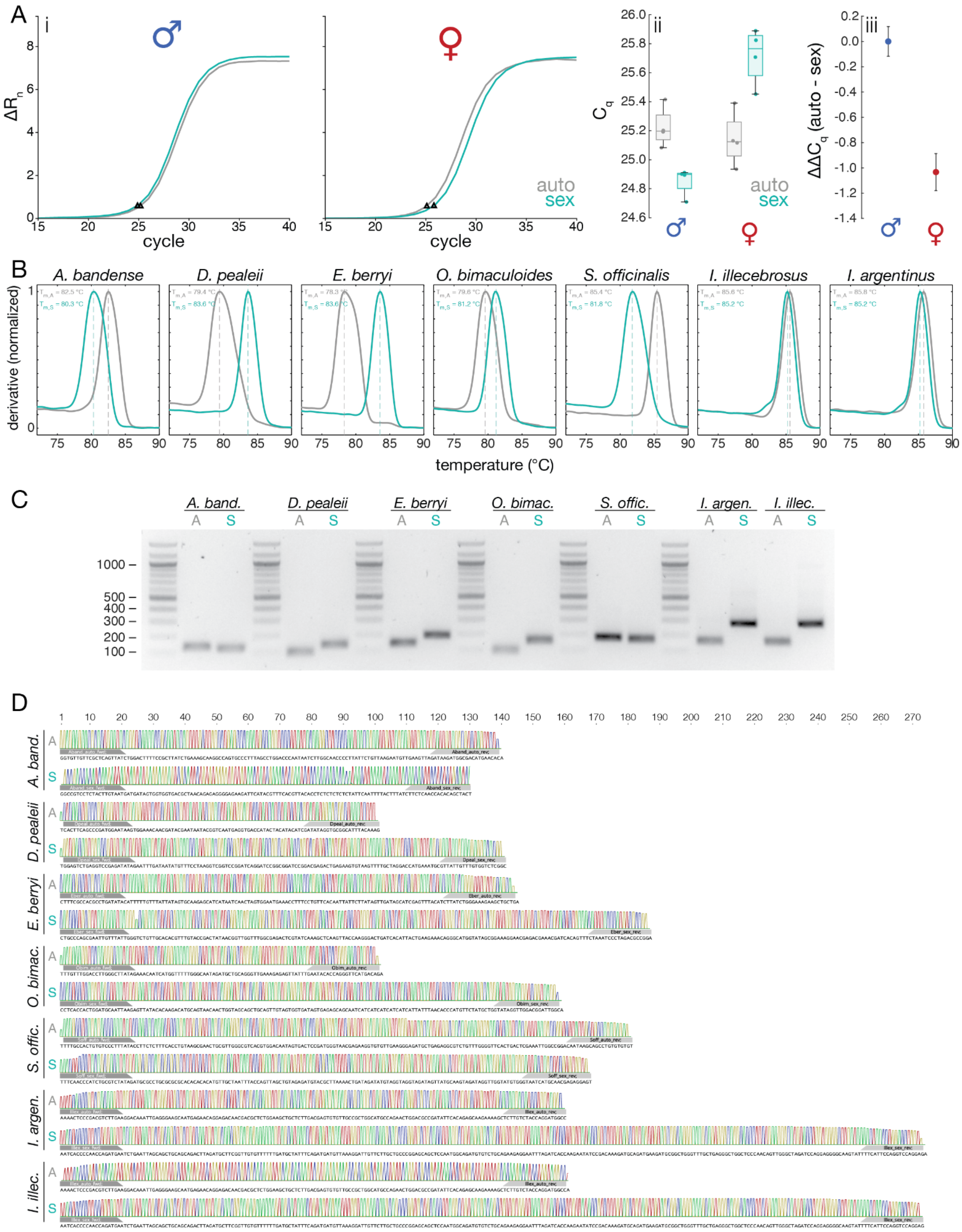


**Figure S2.** Validation of primer pairs used for quantitative PCR against autosome and sex chromosome loci of seven cephalopod species. (A) determination of $\Delta\Delta C_q$ values for a single male and a single female standard of *A. bandense*: (i) amplification curves of male and female standards using autosome- and sex chromosome-targeting primer pairs (specified in Table S1). Triangles (△) represent $C_q$ values as determined by Design & Analysis Software. For visual clarity, only a single technical replicate is shown for each; (ii) $C_q$ values from four technical replicates of each reaction. Box plots show the interquartile range with median line; whiskers indicate minimum and maximum values. (iii) Normalized $\Delta\Delta C_q$ values determined from four technical replicates (see Quantitative PCR analysis in Methods). Error bars indicate the summation of standard error of the mean values of $C_{q,auto}$ and $C_{q,sex}$. (B) Melt curve analysis of all primer pairs used in this study (Table S1). Melting temperature ($T_m$) as determined by Design & Analysis Software is indicated by a dashed line. (C) Agarose gel electrophoresis of qPCR amplicons. (D) Raw signal trace from nanopore sequencing of qPCR amplicons, with primer binding sites indicated.

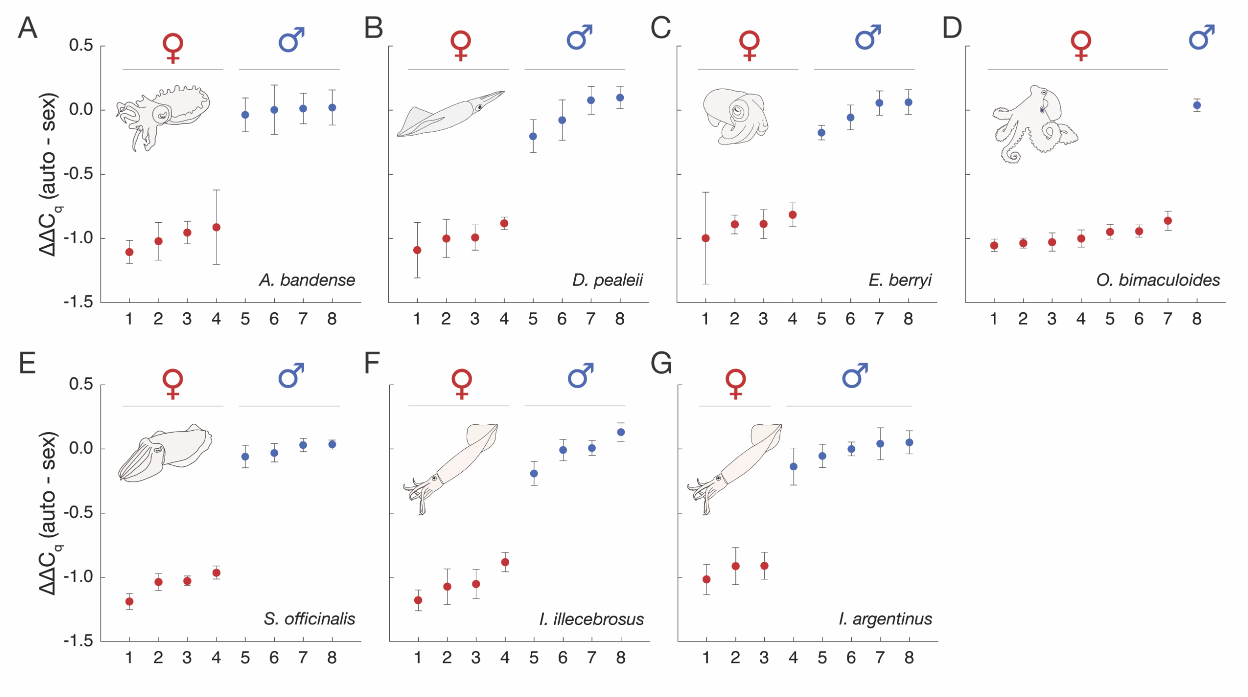


**Figure S3.** (A) *A. bandense.* (B) *D. pealeii.* (C) *E. berryi.* (D) *O. bimaculoides.* (E) *S. officinalis.* (F) *I. illecebrosus.* (G) *I. argentinus.* For each species, $\Delta\Delta C_q$ values were calculated between autosomal and sex-linked loci across eight blinded, randomized tissue samples, normalized such that the median value of the greater-$\Delta\Delta C_q$ cluster (male) was set to zero. Individuals are arranged along the x-axis in order of increasing $\Delta\Delta C_q$. Data points and error bars represent the mean and standard error of the mean, respectively, of four technical replicates.

A

**Autosome**

```
               Illex.1:    1-  100
I. illecebrosus AAAACTCCCGACGTCTTGAAGGACAAATTGAGGGAAGCAATGAGAACAGGAGACAACGACGCTCTGGAAGCTGCTCTTGACGAGTGTGTTGCCGCTGGCA
               ||||||||||||||||||||||||||| |||||||||| ||||||| ||||||||| ||||||||||||| || |||||| || ||| | ||||||||||
     D. pealeii AAAACTCCCGACGTCTTGAAGGACAAACTGAGGGAAGCCATGAGAAGAGGAGACAAGGACGCTCTGGAAGATGTTCTTGATGAATGTATCGCCGCTGGCA
               Dpe01:65236255-65236100:    1-  100

               Illex.1:  101-  160
I. illecebrosus TGCCAGAACTGGACGCCGATATTCACAGAGCAAGAAAAGCTCTTGTCTACCAGGATGGCC
               |||||||| | ||||| |||||||||||||||||||||||||||| | ||||||||||||
     D. pealeii TGCCAGAATTAGACGCTGATATTCACAGAGCAAGAAAAGCTCTTGACGACCAGGATGGCC
               Dpe01:65236255-65236100:  101-  160
```

B

**Sex chromosome**

```
               Illex.Z:1-100
I. illecebrosus AATCACCCCAACCAGATGAATCTGAATTAGCAGCTGCAGCAGACTTAGATGCTTCGGTTGTGTTTTTTGATGCTATTTCAGATGATGTTAAAGGATTGTT
               |||| ||||| |||||||||||||| ||||||||||||||||||||||||||||| |||||||| |||||  ||||| ||||||||  || |||||||||
     D. pealeii AATCCCCCCAGCCAGATGAATCTGAGTTAGCAGCTGCAGCAGACTTAGATGCTTCAGTTGTGTTATTTGAATCTATTGCAGATGATAGTATAGGATTGTT
               Dpe43:25792259-25792529:1-100

               Illex.Z:101-200
I. illecebrosus CTTGCTGCCCGGAGCAGCTCCAATGGCAGATGTGTCTGCAGAAGAGGAATTTAGATCACCAAGAATATCCGACAAAGATGCAGATGAAGATGCGGCTGGG
               ||| ||||| |||||||||||||||||||||||||| |||| |  || ||| |  || ||||  | |||   |||||| ||||||||||||||||||||
     D. pealeii CTTACTGCCTGGAGCAGCTCCAATGGCAGATGTGTCCGCAG-A-TGG-ATTCATCTCGCCAAACAAATCTACCAAAGAGGCAGATGAAGATGCGGCTGGA
               Dpe43:25792259-25792529:101-197

               Illex.Z:201-273
I. illecebrosus TTTGCTGAGGGCTGGCTCCCAACAGTTGGGCTAGATCCAGGAGGGGCAAGTATTTTCATTCCAGGTCCAGGAG
               ||||||||| ||||| |||||||||||||||||||||||||||| ||||||||||||||| ||||||||||||
     D. pealeii TTTGCTGAGTGCTGGATCCCAACAGTTGGGCTAGATCCAGGAGGTGCAAGTATTTTCATTGCAGGTCCAGGAG
               Dpe43:25792259-25792529:198-270
```

**Figure S4.** Amplicons used for sex genotyping in *I. illecebrosus*, showing the alignment of *I. illecebrosus* reads to the *D. pealeii* genome. (A) Autosomal amplicon (aligned to *D. pealeii* chromosome 1). Red line marks the *I. illecebrosus* primers. The forward primer has no mismatches, while the reverse primer has mismatches at position 13 (C>A) and position 15 (T>A). (B) Sex (Z) chromosome amplicon (aligned to *D. pealeii* chromosome 43). Blue line marks the *I. illecebrosus* primers. The forward primer has two mismatches (position 5 C>A and position 11 G>A), while the reverse primer has one mismatch at position 13 (C>G).

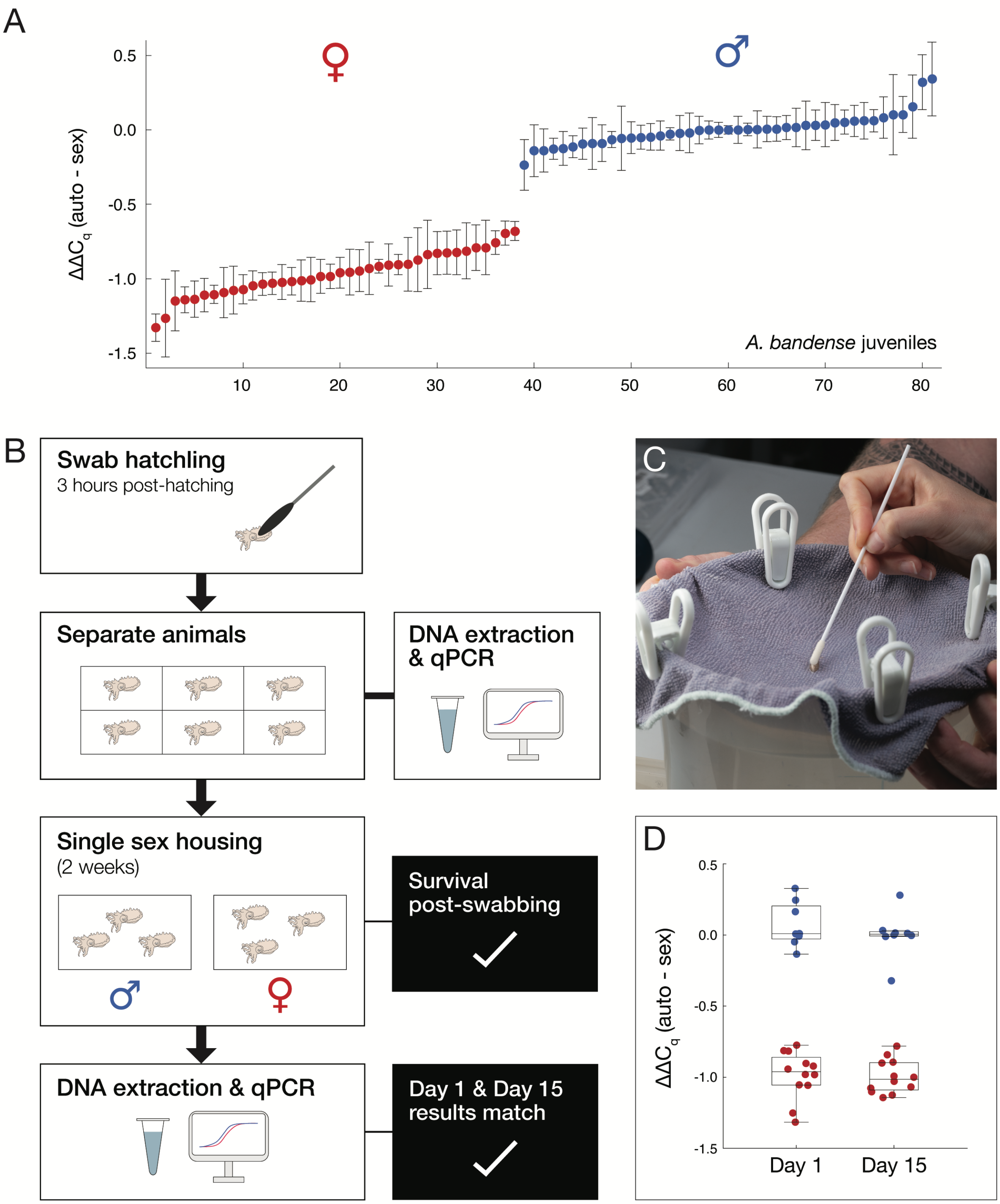


**Figure S5.** (A) Ranked plot of $\Delta\Delta C_q$ values comparing autosomal and sex loci in live juvenile *A. bandense* individuals, normalized such that the median value of the male cluster is set to $\Delta\Delta C_q = 0$. Each data point represents the mean of four technical replicates from a single animal (N = 81 animals), with error bars representing standard error of the mean. Predicted sex based on a $\Delta\Delta C_q$ threshold of -0.5 is shown in red (female) or blue (male); all predictions were confirmed by dissection. (B) Experimental overview: Hatchlings were swabbed within three hours post-hatching (3 hph) and separated into individual tanks.

Genomic DNA was extracted from the skin swab and used for qPCR-based sex prediction. After swabbing, the hatchlings were grouped into single-sex tanks for two weeks to monitor their survival. After two weeks, the animals were euthanized, DNA extracted from tissue, and used as input for qPCR to confirm that skin swabs and tissue extractions show matching sex predictions. (C) To swab hatchlings, the animals are placed on a towel soaking in a jug of seawater. (D) $\Delta\Delta C_q$ values obtained for day-1 (skin swabs) and day-15 (tissue extraction) juveniles. Each data point represents the mean of four technical replicates from a single animal (N = 20 animals). Sex predicted from day-1 samples is shown in red (female) or blue (male) and was used for single-sex housing; all animals assayed on day 15 clustered according to their day-1 prediction, using a cutoff value of $\Delta\Delta C_q$ = -0.5.
